# Automatic Bevacizumab Response Prediction in Ovarian Cancer from Digital Pathology Images via Novel AI-based Computational Pipeline

**DOI:** 10.64898/2026.04.29.721782

**Authors:** Abdullah Alsaiari, Turki Turki, Y-h. Taguchi

**Affiliations:** Department of Computer Science, King Abdulaziz University, Jeddah 21589, Saudi Arabia; Computer Department, Applied College, Najran University, Najran, 66462, Saudi Arabia; Department of Physics, Chuo University, Tokyo 112-8551, Japan

**Keywords:** Gynecological cancer, ovarian cancer, bevacizumab response, histopathology images, deep and machine learning, advanced AI applications in gynecological cancer

## Abstract

Ovarian cancer is a gynecological cancer, which, if metastasized and not detected early, can cause death among women. Therefore, accurate prediction of drug responses to ovarian cancer is needed. A gynecological pathologist inspects abnormality in tissues and provides a report for patients; however, this diagnostic process (1) is difficult to undertake; (2) requires experience; and (3) is time-consuming. Moreover, existing tools are imperfect. Hence, we present a computational pipeline to improve predictions of drug response pertaining to ovarian cancer. First, we downloaded digital pathology images pertaining to ovarian responses to bevacizumab from the Cancer Imaging Archive Repository. We employed a histogram of oriented gradients for images, constructed feature vectors, and used Fisher’s linear discriminant analysis to alter data representations through dimensionality reduction. This reduced-dimensionality data was used for regression analysis, employing support vector regression coupled with various kernels and calculating the area under the ROC curve (AUC). Experimental results were validated using transformer-based models (ViT and Swin) and other deep learning (DL) models (VGG16, ResNet50, InceptionV3, MobileNetV2, and EfficientNetB6). Our approach using a radial kernel (named SVRD+R) improved AUC performance by 17% compared to the best-performing transformer-based model (ViT). Likewise, AUC performance improved by 14.9% when compared against the best DL-based model (MobileNetV2). These results demonstrate feasibility, showing that induced models via the presented AI-based pipeline can lead to superior performance when investigating prediction problems pertaining to gynecologic cancer studies.

**MSC:** 92B05; 68T09

## 1. Introduction

Ovarian cancer constitutes one of the leading causes of death related to gynecological cancer worldwide, mainly due to the high rates of diagnosis at advanced stages and the substantial variations in treatment response [5, 6]. Thus, accurate assessment of treatment outcomes is crucial for guiding treatment strategies and enhancing long-term survival [7]. Over the past few years, digital pathology has promoted the quantitative extraction of biomarkers from high-resolution histopathological images to guide data-driven approaches to precision oncology [8]. Despite this, conventional diagnostic workflows still rely heavily on manual visual assessment by pathologists, which is time-consuming and subject to intra- and inter-observer variability [9].

Existing studies have aimed to tackle this binary classification task using clinical, genomic, and histopathological images [11,12,13,14,15,16]. However, performance results for predicting drug response to ovarian cancer using histopathological images are imperfect, and an advanced AI-driven approach applied to larger sample sizes is needed [1, 2]. Such an approach requires validation on the whole dataset. Therefore, we present a novel AI-based computational approach to improve drug response in ovarian cancer. We evaluate our model against deep learning and transformer-based models. In total, we explore seven randomly selected baseline methods, including five convolutional neural networks, namely Visual Geometry Group-16 (VGG16), Residual Network-50 (ResNet50), InceptionV3 (MobileNetV2, and EfficientNetB6), and two transformer-based models (Vision Transformer (ViT) and shifted windows (Swin)). Each induced model was finely tuned to identify effective and ineffective (invalid) treatment responses. Models induced via our AI-based pipeline employ the following feature extraction methods: histograms of oriented gradients (HOG) and local Fisher’s discriminant analysis (LFDA). Constructed feature vectors are then coupled with support vector regression (SVR), and the four kernel functions (radial, linear, polynomial, and sigmoid). The results demonstrate the superior performance of our models.

The key contributions of this study are the following:

1. We present an AI-based pipeline that incorporates a computational trick (turning classification into a regression problem to solve the classification problem) composed of feature extraction and dimensionality reduction via HOG and LFDA. We then employ supervised regression training (rather than classification) using support vector regression (SVR). Four induced regression models (SVRD+L, SVRD+P, SVRD+R, SVRD+S) were attributed to four kernels (linear (L), polynomial (P), radial (R), and sigmoid (s)) and compared against seven deep and transformer-based models, including VGG16, ResNet50, InceptionV3, MobileNetV2, EfficientNetB6, Swin, and ViT.
2. The best performance setting was identified in our pipeline when coupled with SVRD+R by searching a dimensionality reduction feature range of two to fifty-four. Experimental results demonstrate that the SVR with a radial basis kernel and 48-dimensional feature vectors from LFDA outperformed all evaluated deep learning models.
3. The results, after performing five-fold cross-validation on the complete dataset of 250 histopathology images, demonstrate that our model, SVRD+R, outperforms the best transformer-based model, ViT, with a 17% improvement in AUC performance. Moreover, we outperformed the best deep learning-based model, MobileNetV2, with a 14.9% improvement in AUC performance. These results provide a systematic comparison of deep learning and traditional machine learning strategies in digital pathology and establish a potential framework for future research on predicting treatment responses in ovarian cancer.

## 2. Literature Review

To improve the objectivity, scalability, and clinical applicability of oncology decisionmaking, recent efforts have focused on developing automated and repeatable computational processes that capture subtle morphological patterns associated with treatment response [10]. Mallya et al. [20] experimented with various deep learning-based models to predict bevacizumab response in ovarian cancer patients using histopathological whole-section images from the cancer imaging archive (TCIA) database. An analysis of 287 H&E (hematoxylin and eosin)-stained whole slide images from 78 patients was performed to determine whether treatment was effective or ineffective. They evaluated several deep learning models, including ResNet50, Dense Network-121 (DenseNet121), ViT, and Swin, to name a few. The deep learning-based model UNI+ VarMIL combines models with multi-instance learning (MIL) and was trained on 212 labeled whole slide images (WSIs). This resulted in a balanced accuracy of 69.8% when tested on only 74 WSIs of uniform class distribution. Liu et al. [21] proposed a machine learning-based approach to predict prognosis and drug response in ovarian cancer that works as follows. First, they employed bioinformatics-based analysis using weighted gene co-expression network analysis (WGCNA) and CIBERSORT, applied to RNA-seq data and genomic mutations in ovarian cancer, thereby identifying 1582 macrophage M2-related genes. These 1582 genes were then subjected to univariate Cox analysis, yielding 34 prognostic biomarkers. Datasets from the Gene Expression Omnibus (GEO) with access numbers GSE14764 and GSE140082 were used to validate the discriminative power of these 34 biomarkers. Machine learning models integrating various learning algorithms were employed on these datasets with 34 biomarkers, constructing macrophage eM2-related gene signatures. Based on multiple external validation cohorts, the resulting model achieved a consensus index of about 0.68-0.75.

Ahn et al. [22] demonstrated that a deep learning-based histopathology classifier, PathoRiCH, could be used to predict high-grade serous ovarian cancer’s response to platinum-based therapy. According to their study, 814 patients from three cohorts (SEV, TCGA, and SMC) were included in the study, and the classification task was modeled as binary based on a platinum-free interval for each cohort. Training was completed using 754 WSIs from 394 patients, while the other 516 and 136 WSIs were obtained from the remaining patients. As a result of employing multi-instance learning (MIL) across multiple tissue representations (here, cancer tissue was segmented in all tissues), along with multiple magnifications (5x, 20x, and multi-scale), several MIL model variants were produced. Among them, the 20× cancer segmentation model performed best, with an AUC-ROC of around 0.602. Hence, the authors found that combining PathoRiCH prediction with biomarkers such as BRCA and HRD status improved patient risk stratification. Yang et al. [23] developed a graph-based deep learning framework called OCDPI (Ovarian Cancer Digital Pathology Index) that uses H&E-stained whole-section images to predict ovarian cancer prognosis. The model combines a transformer-based feature extraction technique (CTransPath [27]) with a graph neural network for prognosis prediction. The OCDPI performed well in the two external validation cohorts. They obtained a hazard ratio (HR) of 1.916 with a 95% confidence interval (CI) of 1.38-2.66 in one of the cohorts, while obtaining an HR of 2.796 with a 95% CI of 1.40-5.56 for the other cohort.

Zhang et al. [24] developed a deep learning model (KANSurv) for subtype identification of ovarian cancer patients with different drug responses. Their model was trained on gene expression data of 360 ovarian cancer patients from TCGA. The model was then tested on 415 samples of gene expression data from the Gene Expression Omnibus (GEO) with accession numbers GSE51088, GSE17260, and GSE26712. Compared to other models, their model performance ranged from 0.60 to 0.63 when the C-index was considered.

Crispin-Ortuzar et al. [25] proposed a machine learning framework (IRON) to predict neoadjuvant chemotherapy response to high-grade serous ovarian cancer as follows. First, they processed clinical, genomic, and radiomic data, constructing feature vectors and feature selection. Then, constructed feature vectors were provided to train an ensemble of machine learning algorithms (elastic net, support vector regression, and random forest) on 72 samples. Then, prediction was performed on 42 samples and an AUC of 0.80 was obtained. Mendes et al. [26] proposed a machine learning approach to identify discriminative metabolic features between high-responders (HRs) and low-responders (LRs), followed by evaluations of the performance. The models included partial least-squares discriminant analysis (PLS-DA), sparse PLS-DA (sPLS-DA), and random forests (RFs), applied to healthy controls and (carboplatin + paclitaxel) ovarian cancer patients treated with drug combinations. PLS-DA achieved an AUC of 0.98, discriminating between HRs and LRs when nine samples were utilized.

**Table 1.** Summary of related studies for treatment response and prognosis prediction in ovarian cancer. OV is ovarian cancer. TCGA is the Cancer Genome Atlas. OCDPI is the ovarian cancer digital pathology index. HMUCH is the Harbin Medical University Cancer Hospital. PLCO is a prostate, lung, colorectal, and ovarian cancer screening trial. WSIs are whole-slide images. HOG is the histogram of oriented gradients. LFDA is local Fisher’s discriminant analysis. AUC is the area under the curve.

| Year | Study | Data source | No. of models | Best model | Class | Testing | Results |
| --- | --- | --- | --- | --- | --- | --- | --- |
| 2022 | Mendes et al. [26] | Ex vivo ovarian tumor tissue cultures (patient-derived explants) | 1 | PLS-DA (20-feature biomarker panel) | 2 class (high responder vs low responder) | 9 patient-derived tumor samples | AUC = 0.98 (Accuracy = 90%) |
| 2023 | Crispin-Ortuzar et al. [25] | Baseline CT scans + clinical data (NeOV cohort for training/hold-out and Barts cohort for external validation) | 1 | IRON full model (clinical + CA-125 + radiomics + ctDNA) | 2 class (responder vs stable/progressive; RECIST 1.1) | 42 CT scans (external validation cohort—Barts) | AUC = 0.80 |
| 2024 | Yang et al. [23] | TCGA-OV (WSIs), PLCO cohort, HNUCH cohort | 1 | OCDPI (Graph DL + Transformer) | 2 class (low OCDPI vs high OCDPI) | 608 WSIs (PLCO); 94 WSIs (HNUCH) | HR = 1.916 (PLCO); HR = 2.796 (HNUCH); log-rank P < 0.01 |
| 2024 | Ahn et al. [22] | TCGA-OV, SEV cohort, SMC cohort | 6 | PathoRiCH (cancer-segmented area, 20× MIL) | 2 class (favorable vs poor response to platinum) | 516 WSIs (TCGA external validation set) | AUC = 0.602 |
| 2024 | Zhang et al. [24] | TCGA-OV, GEO datasets (GSE17260, GSE26712, GSE51088) | 5 | KANSurv (DeepSurv + Kolmogorov–Arnold Network) | subtype identification of ovarian cancer patients with different drug responses. | 415 gene expression samples from (GSE51088, GSE17260, and GSE26712) | C-index ≈ 0.60–0.63 |
| 2025 | Mallya et al. [20] | TCIA (WSIs) | 13 | UNI + MIL | 2 class (responder/non-responder) | 74 WSIs | Balanced ACC (AUC) = 69.8% |
| 2025 | Liu et al. [21] | TCGA and GEO | 52 | LASSO-based ML signature | 2 class (high-risk vs. low-risk patients) | Gene expression data (GSE14764 and GSE140082) | C-index ≈ 0.68–0.75 |
| 2026 | Proposed | TCIA (WSIs) | 11 | HOG + LFDA +<br>SVR (RBF), named<br>SVRD+R | 2 class (effective vs<br>invalid) | 250 WSIs | AUC = 0.7854 |

## 3. Materials and Methods

### 3.1 Dataset Description

This study was conducted using data from the Cancer Imaging Archive (TCIA) at https://www.cancerimagingarchive.net/collection/ovarian-bevacizumab-response/. In the original dataset, there are 287 whole slide images (WSIs) of ovarian cancer that contain information about treatment response for each image. The class distribution includes 160 WSIs categorized as “effective” when ovarian cancer responds to the bevacizumab treatment, and 127 WSIs categorized as “invalid” when ovarian cancer does not respond to the treatment. These labeled images served as the gold standard throughout the study. To prepare the whole-slice images, preliminary preprocessing is required since the images are extremely large and contain heterogeneous regions (including tumor tissue, stroma, necrotic zones, and background). During this stage, all slides that contain duplicate recordings, unreadable areas, or inadequate tissue content are excluded. Further processing of the remaining WSIs was then conducted, and representative tissue regions suitable for subsequent calculations and analyses were selected, and representative tissue regions suitable for subsequent calculations and analyses were extracted. The final dataset contains a random selection of 250 high-quality JPEG images divided into 125 effective and 125 invalid samples after preprocessing and extraction. Then, in all machine learning and deep learning experiments, this image dataset was consistently used as an input for feature extraction, dimensionality reduction, and classification.

### 3.2 Local Fisher Discriminant Analysis (LFDA)

Following feature extraction using a histogram of oriented gradients (HOGs) [17], local Fisher’s discriminant analysis (LFDA) was used to reduce data dimensionality and enhance class separation while maintaining local structure. LFDA is an extension of Fisher’s linear discriminant analysis (LDA), which is useful for analyzing complex and potentially multimodal data distributions that are commonly encountered in histopathological imaging analyses [3,4].

Given a training set 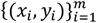 in which *x*_*i*_ ∈ ℝ^*n*^ denotes the input feature vector and *y*_*i*_ represents the corresponding class label. LFDA aims to learn a linear projection matrix *W* ∈ ℝ^*n*×*r*^. It projects the original feature space onto a lower-dimensional subspace while preserving locally discriminative information. LFDA aims to maximize the ratio between the scatter between classes and the scatter within classes. The selected projection matrix is obtained by solving the following optimization problem:

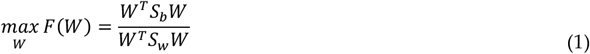

where *S*_*b*_ indicates the between-class scatter matrix and *S*_*w*_ corresponds the within-class scatter matrix. LFDA incorporates locality information by weighing neighboring samples, allowing it to better capture local class structures and handle multimodal class distributions. We projected data (i.e., feature vectors with ℝ^*m*×*n*^) onto *W* with ℝ^*n*×*r*^ to obtain data with reduced feature space (i.e., ℝ^*m*×*r*^), where *r*<<*n*. In this study, LFDA was applied to reduce the dimensionality of HOG features across a range of reduced dimensions prior to classification using support vector regression (SVR), enabling an effective balance between discriminative power and feature compactness for small-to-medium-sized histopathology datasets.

### 3.3 ε-Support Vector Regression (SVR)

In this study, support vector regression (SVR) is used as a kernel function-based learning method to model potential nonlinear relationships. Let the training dataset be denoted as 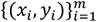 in which *x*_*i*_ ∈ ℝ^*n*^ represents the feature vector of the *i*-th sample and *y*_*i*_ ∈ ℝ denotes the corresponding label. To enable nonlinear modeling, SVR was formulated in its dual form by introducing Lagrange multipliers *α*_*i*_and *α*^*^. Here, *ε* defines the tube around the regression line where errors inside the tube are ignored (i.e., not penalized). *C* is a regularization parameter, which controls overfitting. The resulting dual optimization problem is given in Equation (2) [18] as follows:

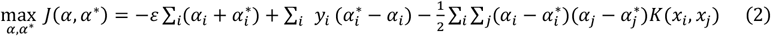

which is subject to

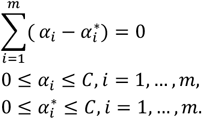

From the optimal solution of Equation (2), the weight of genes is recovered as follows:

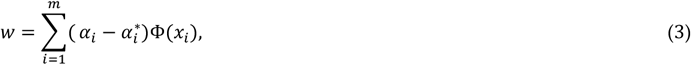

where Φ(*x*_*i*_) denotes the nonlinear mapping of the input sample into the feature space induced by the selected kernel function.

Accordingly, the regression function used to predict a new sample *x*′ is expressed as follows:

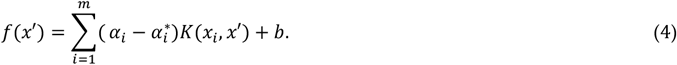

Four kernel functions are considered to capture different types of nonlinear relationships in the feature space: linear kernels, polynomial kernels, radial basis functions (RBFs), and sigmoid kernels. Using a high-dimensional feature space, the SVR model can establish linear or nonlinear decision boundaries by defining different kernel functions for similar measures between input samples [19]. The mathematical formulations of the employed kernel functions are given below [18]:

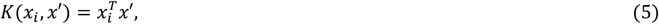

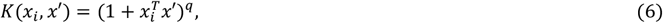

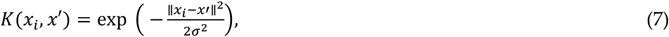

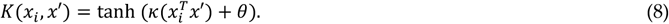

There are four types of kernels used in the analysis: the linear kernel calculates the inner product between feature vectors, the polynomial kernel models the interaction of higher-order features, and the radial basis function (RBF) kernel measures similarity based on Euclidean distances between the samples. The sigmoid kernel has a nonlinear mapping that is similar to neural activation functions. It is evident that our goal is to induce a repression model *f*: *x*′ → *y*^′^, where *y*^′^ ∈ ℝ that is comparable to using AUC against induced classification models *h*: *x*′ → *y*^′′^, where *y*^′′^ ∈ {invalid, effective}. Therefore, Equation 4 for prediction can be written as follows:

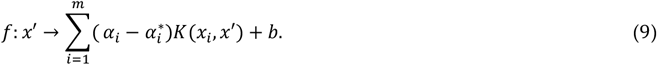

Figure 1 illustrates how the proposed pipeline for predicting ovarian cancer treatment response works. There are two main parts: data preparation and the utilization of artificial intelligence-based methods. The former is used to collect data, and the latter constructs feature vectors and builds models. For data preparation, ovarian cancer specimens are removed by the treating gynecologist, followed by a pathologist who aids in producing digitized WSIs. Then, the original svs images of 54342 × 41048 pixels are resized to average JPEG images of 256 × 256 pixels. Unlike existing studies that generally aim to employ deep and machine learning techniques under the classification setting, we constructed feature vectors from these JPEG images using HOG and FLDA, which were provided to train SVR under the regression setting. Then, the class labels of unseen histopathology images were predicted, followed by evaluation using AUC, as described in the following sections.

**Figure 1.**
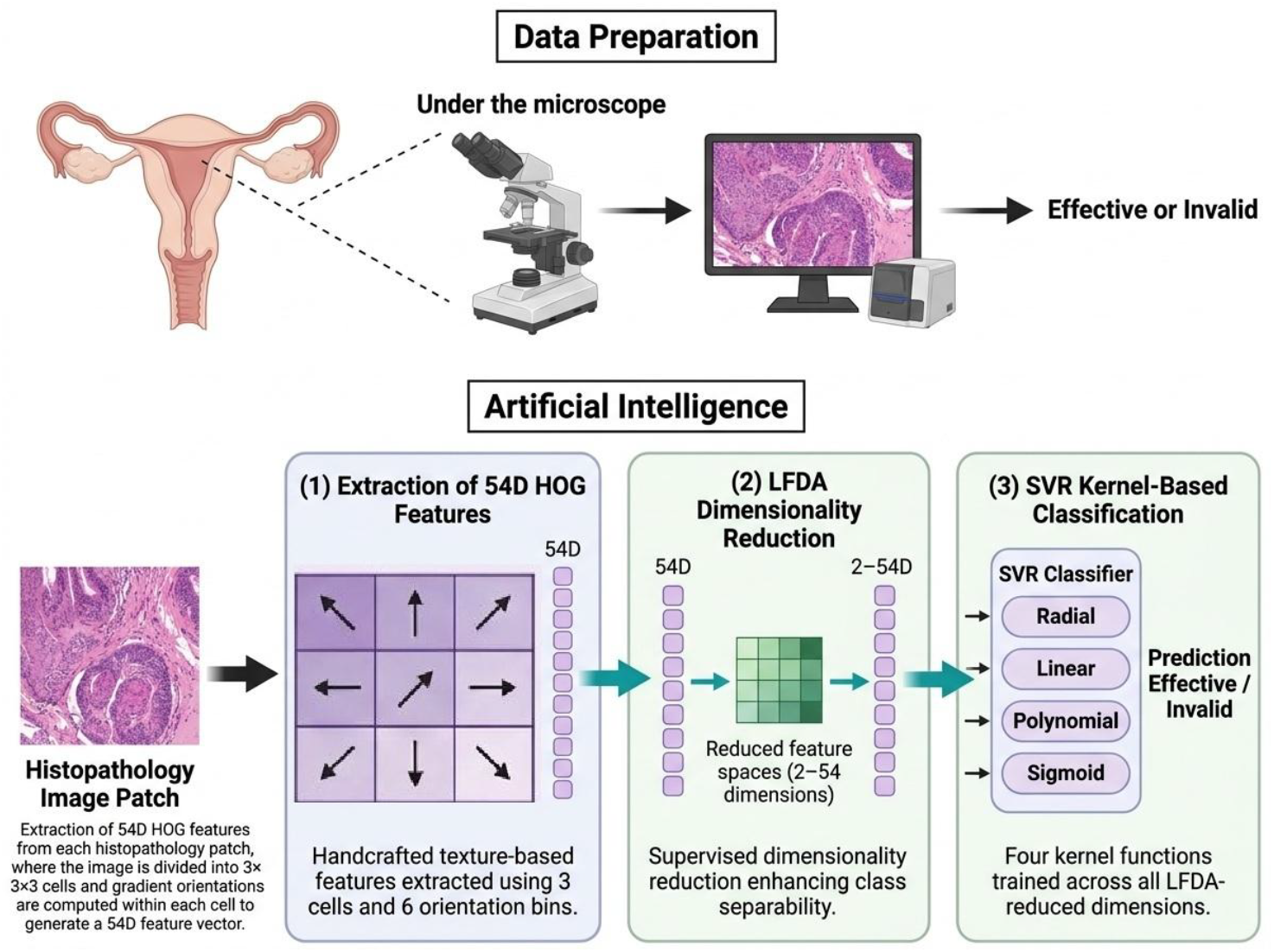
Overview of the proposed AI-based pipeline. Data Preparation: Histopathology images of ovarian cancer response to bevacizumab treatment, obtained from the Cancer Imaging Archive. Artificial Intelligence: HOG was applied to extract features of dimensions from 2 to 54, followed by further dimensionality reduction using LFDA. Then, data from this final representation were provided to SVR with different kernels, resulting in four models, namely SVRD+L, SVRD+P, SVRD+R, and SVRD+S.

## 4. Experiments and Results

### 4.1 Deep Learning Results

Seven deep learning architectures, including transformer-based models, were randomly selected and used in the experiments: VGG16, ResNet50, InceptionV3, MobileNetV2, EfficientNetB6, Swin Transformer (Swin), and Vision Transformer (ViT). All models were trained on the same carefully selected dataset and under the same training process to ensure a fair comparison.

As shown in Figure 2, the training accuracy and loss of all deep and transformer-based models over ten training epochs are plotted. Transformer-based models tend to converge faster during the initial training epochs, while other deep learning architectures, such as MobileNetV2, converge slowly, achieving stable performance. This behavior can be seen in the loss curves. Compared to deep learning and transformer-based models, MobileNetV2 exhibits more stable loss patterns across training epochs, as shown by the progressive performance differences.

**Figure 2.**
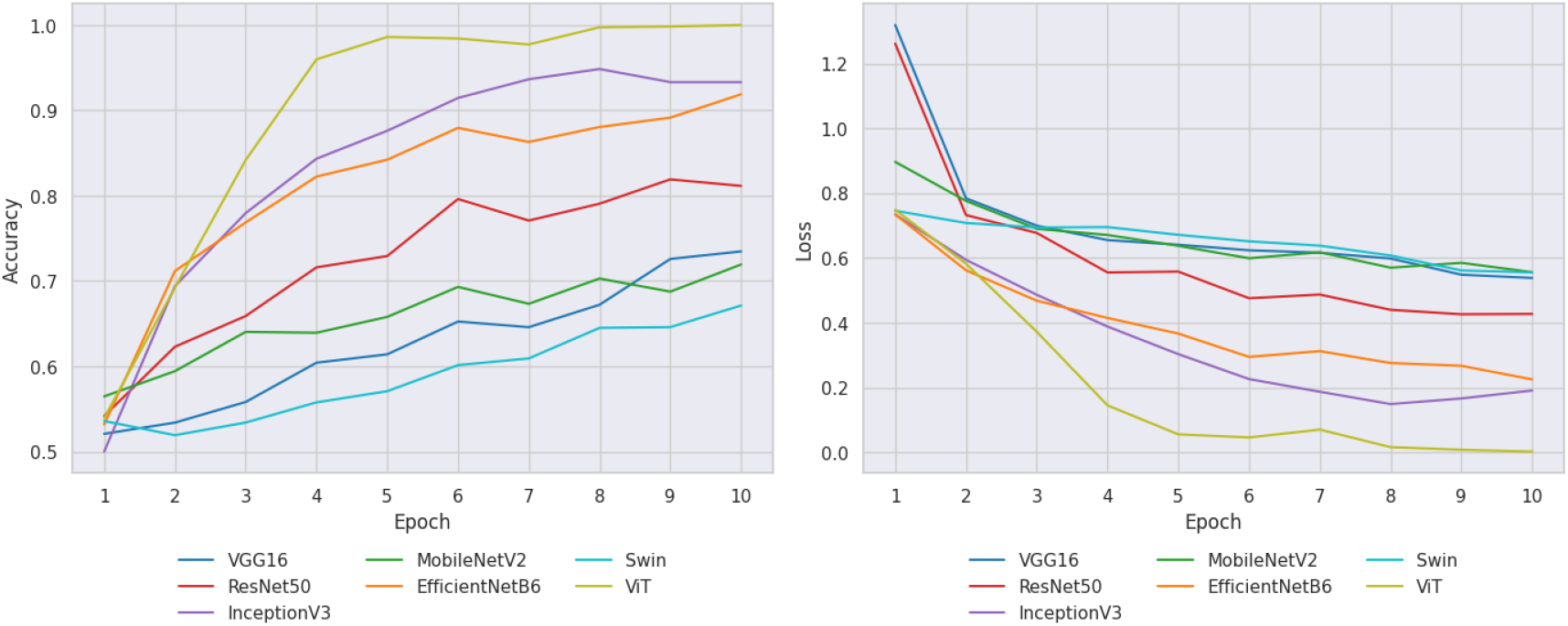
Training accuracy and loss curves of seven deep learning and transformer-based models (VGG16, ResNet50, InceptionV3, MobileNetV2, EfficientNetB6, Swin, and ViT) over 10 training epochs.

Among all the tested architectures in Figure 3, EfficientNetB6 has the highest AUC of 0.665, indicating its relatively good discriminative ability. MobileNetV2 and VGG16 yield similar performance results (AUC =0.613). ResNet50 obtained a marginal performance difference when compared to VGG16 and MobileNetV2. The overall classification performance of convolutional neural networks is moderate, whereas the performance of transformers is significantly lower. However, all deep learning methods have low AUC values, creating a need to develop advanced AI-based pipelines.

**Figure 3.**
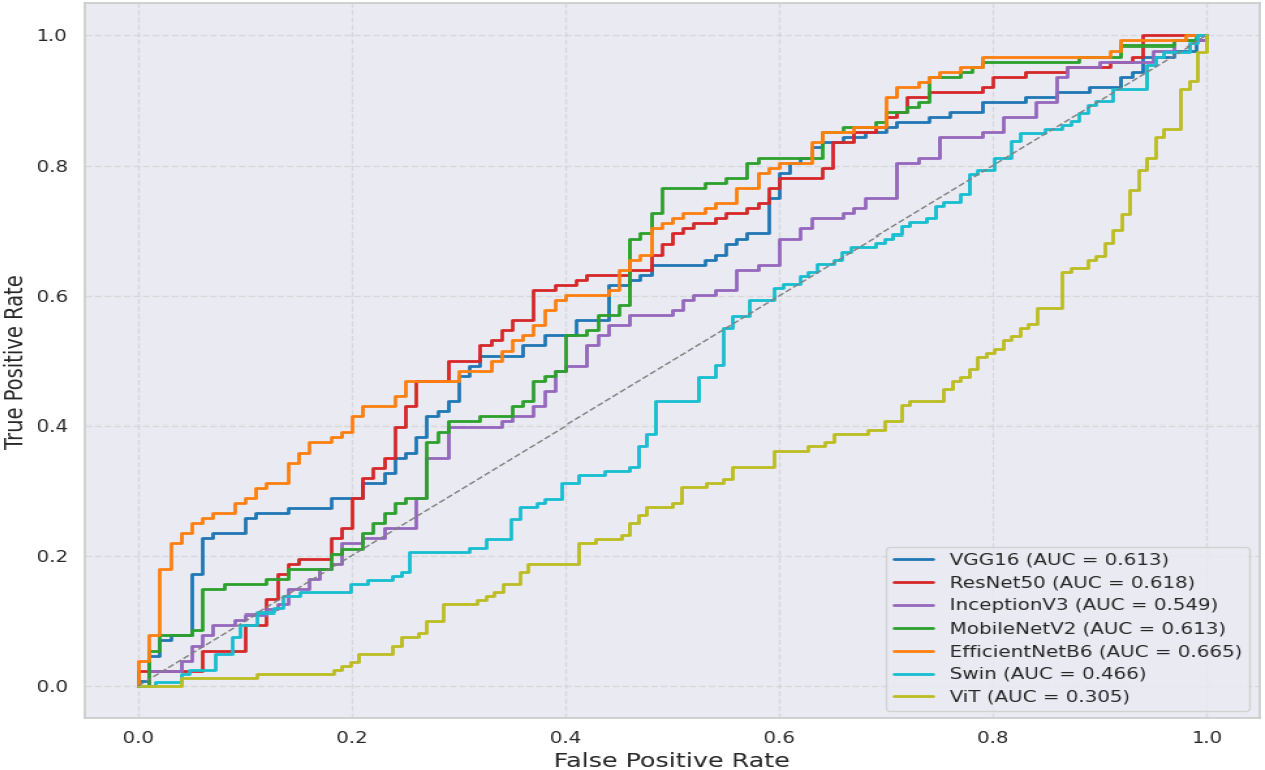
Receiver operating characteristic (ROC) curves and corresponding AUC values for all deep learning and transformer-based models evaluated on the ovarian cancer dataset.

#### 4.2 AI-based Pipeline Results

In machine learning experiments, support vector regression (SVR) is used in our pipeline along with dimensionality reduction features. Our study evaluated the performance of four kernel functions—linear kernel, polynomial kernel (degree = 2), radial basis function kernel, and sigmoid kernel—across a dimensionality reduction feature dimension range of 2 to 54. Model performance was analyzed by measuring the area under the ROC curve (AUC). As can be seen in Figure 4, the average AUC is displayed for each kernel function dimension combination.

**Figure 4.**
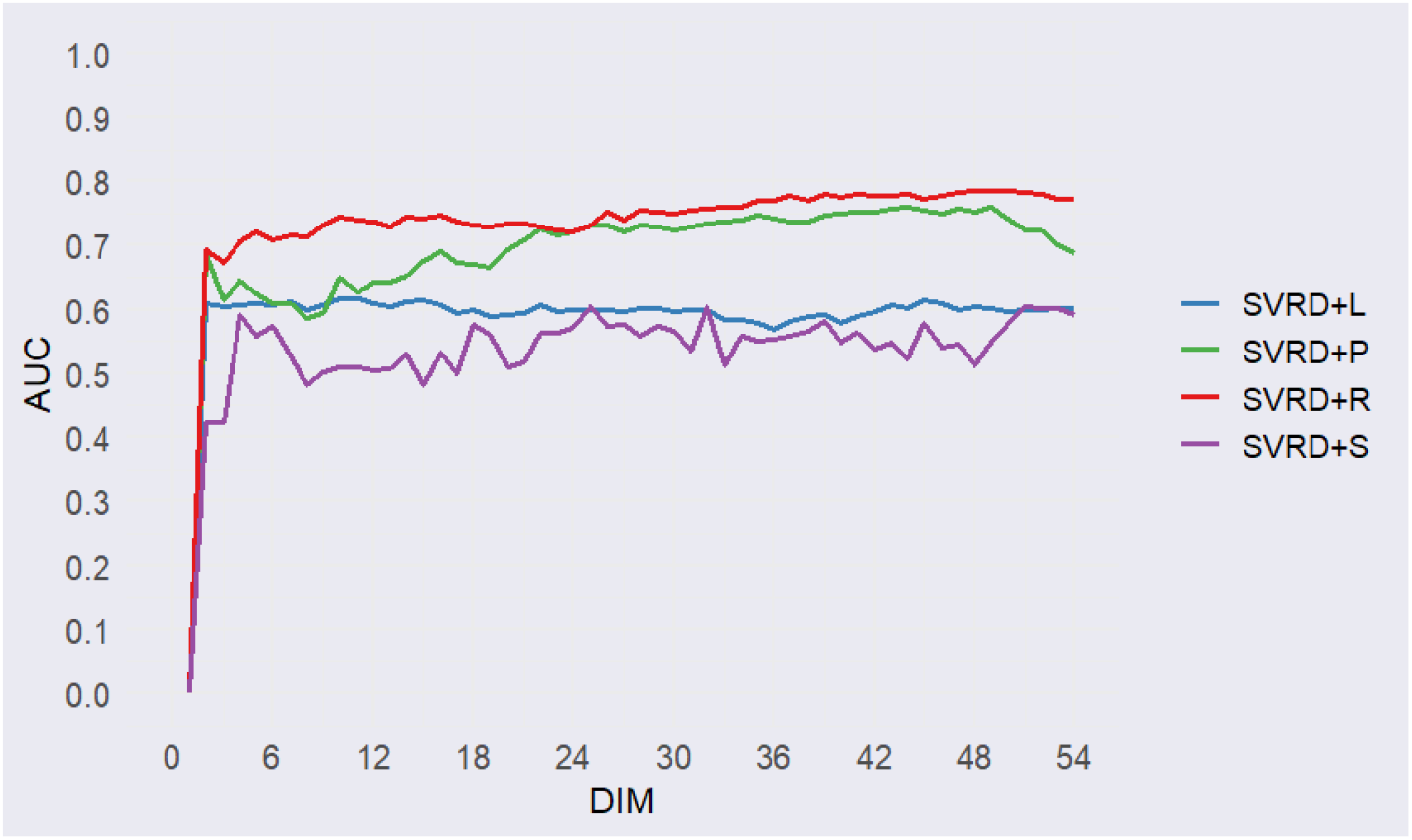
Average AUC performance of models SVRD+L, SVRD+P, SVRD+R, and SVRD+S when coupled with linear, polynomial, radial, and sigmoid kernel functions across different LFDA-reduced feature dimensions (2–54).

As shown in Figure 4, the SVR model using the radial basis function kernel consistently achieved the highest average AUC across the most dimensionality-reduced feature dimensions. The polynomial kernel also demonstrated comparable performance; however, this was slightly inferior to the radial basis function kernel. In contrast, the linear kernel and the sigmoid kernel exhibited significantly lower and less stable discriminative performance across the evaluated dimensionality range. These results indicate that the radial basis function kernel is the most effective configuration for capturing latent patterns related to treatment response prediction in the dimensionality-reduced feature space.

### 4.3 Comparative Analysis Between Deep Learning and AI-based Pipeline Models

To compare deep learning and transformer-based models utilizing five-fold crossvalidation, our pipeline employed a machine learning algorithm (SVR), which exhibited a significant average performance when combined with HOG and LFDA dimensionality reduction. While other models, including VGG16, ResNet50, InceptionV3, MobileNetV2, EfficientNetB6, ViT, and Swin, aim to learn discriminative visual patterns from ovarian cancer image patches, their best performance on the test sets is still inferior by comparison. The best-performing deep learning architecture is MobileNetV2, with an average AUC of approximately 0.636, while the remaining CNN and transformer models exhibit significant fluctuations in performance. In some cases, performance decreases significantly within testing folds. Among the four SVR kernel functions included (radial basis function (RBF), linear kernel, polynomial kernel, and sigmoid kernel), the RBF kernel achieved the highest overall performance when utilized in our pipeline, with an average AUC of 0.785 in 48 dimensions, significantly outperforming the best deep learning model with AUC performance improvements of 14.9%. The polynomial kernel function also performed well (with an AUC of 0.760 in 44 dimensions), confirming the effectiveness of our pipeline in capturing discriminative low-dimensional features extracted from histopathological images. Both the linear kernel function and sigmoid kernel function yielded lower peak AUC values (0.617 and 0.605, respectively); however, both maintained a performance level comparable to that of many deep learning models. This study suggests that dimensionality reduction techniques can enhance class separability in small-to-medium-sized medical image datasets and outperform deep learning architectures. This study shows the highest prediction performance for RBF-SVR, which provides a benchmark for evaluating future improvements.

Figure 5 highlights the differences in AUC performance between deep learning architectures and models induced via our pipeline. SVRD+R and SVRD+P achieved the highest AUC results when compared against deep learning architectures. The results show that training under the regression setting in our pipeline can improve the generalization performance when compared against baseline methods trained under the classification setting, following the typical approach used in previous studies when predicting drug response in ovarian cancer using WSIs.

**Figure 5:**
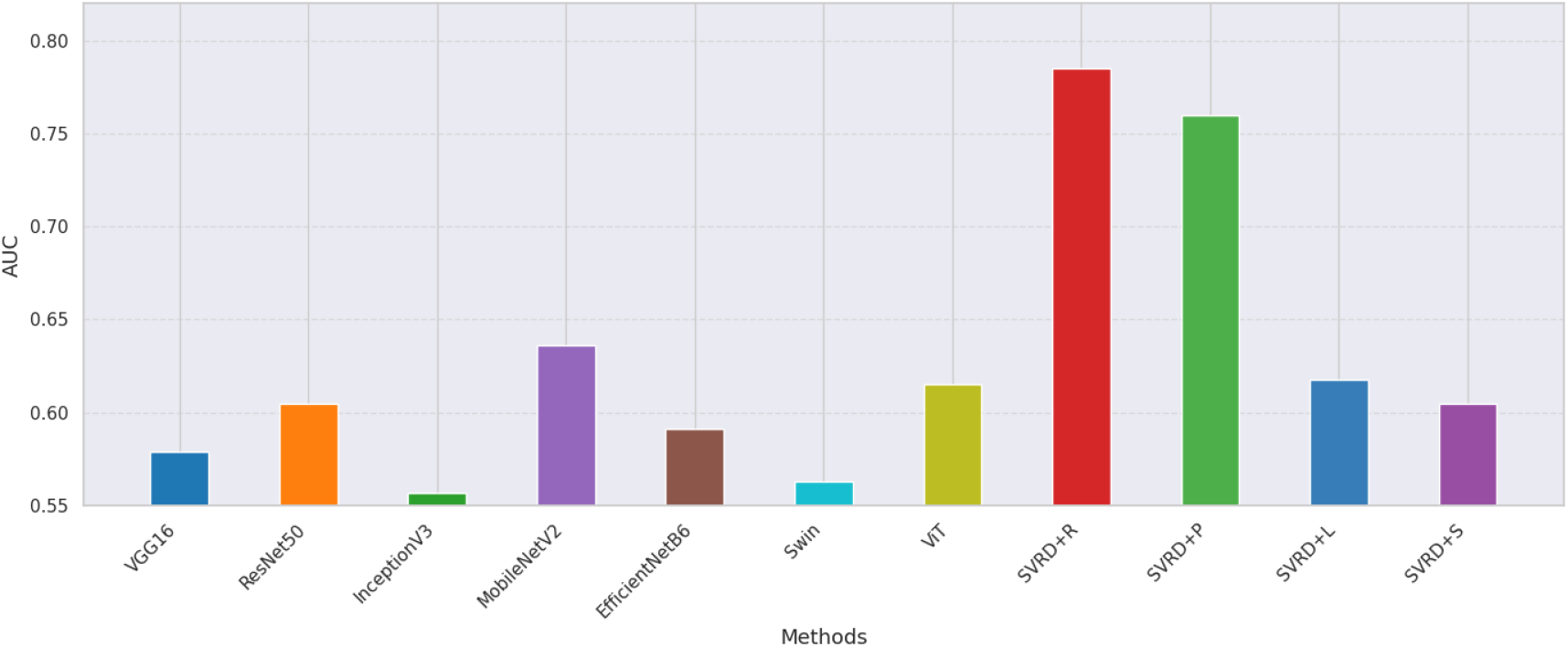
AUC performance results for deep learning architectures and transformer-based models against models SVRD+R, SVRD+P, SVRD+L, SVRD+S based on the complete dataset using five-fold cross-validation.

## 5. Discussion

We processed and balanced the class distribution of 250 histopathological images pertaining to patients with ovarian cancer who were responsive or resistant to the bevacizumab drug. Our proposed AI-based computational pipeline constructs feature vectors of new presentation from histopathological images via incorporating HOG feature extraction, followed by Fisher’s linear discriminant analysis. Data with new representations are then provided to support vector regression (SVR) models with several kernels, including linear, polynomial, radial, and sigmoid. Results based on the complete dataset of 250 histopathological images show that our model SVRD+R yields the best improvements in performance, with a lower bound of 14.9% when the AUC performance metric is considered.

A significant difference in performance was found between the various deep learning models when discriminative information was extracted from histological images of ovarian cancer. As a result of the five-fold cross-validation results, MobileNetV2 demonstrated the best generalization ability and exhibited the lowest variability. The degradation of these DL-based models can be attributed to the fact that small-to-medium-sized datasets are challenging in medical imaging research. In comparison, transformer-based models (including Swin Transformer and Vision Transformer) produced inferior results. The improvement in results produced by our pipeline demonstrates the need for advanced AI-based computational methods to improve performance results.

Compared with deep learning and transformer-based models, our pipeline incorporates a computational trick where the training set was treated as a regression problem. We employed support vector regression (SVR), and data representation was obtained with the help of HOG feature extraction, followed by LFDA dimensionality reduction. Then, a prediction was performed for unseen histopathology images. The results show that when the radial basis function (RBF) kernel function is coupled with SVR, the highest AUC value was obtained among the evaluated kernel functions. These results demonstrate that, when training datasets are small, it is the transfer of models to the problem and the feature extraction method used, combined with appropriate dimensionality reduction, that plays a dominant role in targeting the original classification problem.

It is also important to note that the dimensionality of features has a significant influence on model performance. LFDA results indicate that SVR classifiers perform differently across 2-54 dimensions. This indicates that the features retained greatly affect classification performance. However, despite this difference, the RBF kernel exhibits relative advantage and stability over a wider range of dimensions, providing evidence that it can detect non-linearities in feature extraction. A key consideration in classic machine learning workflows for ovarian cancer prediction is choosing the appropriate level of dimensionality. Selecting an appropriate feature compression level helps balance predictive performance and computational efficiency, especially when datasets are small. It is worth noting that choosing the right kernel (i.e., RBF kernel) in this study was performed using try-and-see manner. Moreover, the weights of pre-trained deep learning and transformer-based models were transferred to the feature extraction part. At the same time, weights in the classification layers were fine-tuned and optimized for the task of predicting bevacizumab drug response.

For experiments run using R, we employed the e1071 package in R for SVR using the svm method [28]. We used the HOG method in the OpenImageR package to run HOG [29], and utilized the LDFA method in the LFDA package to run LFDA [30]. For deep learning and transformer-based models in Python, we employed VGG16 [31], ResNet50 [32], InceptionV3 [33], MobileNetV2 [34], EfficientNetB6 [35], Swin [36], and ViT [37]. We aimed to use well-known histopathological image datasets for the production of bevacizumab drug response in ovarian cancer patients. Such a dataset was provided by Wang et al. [38, 39] from TCIA.

Predicting bevacizumab drug responses in ovarian cancer patients using histopathology images requires a specialized gynecological pathologist. Therefore, the clinical validation and availability of such datasets are not easy. Also, mislabeling results can occur during manual labeling. These factors contribute to the limitations of this study. Experimental results using five-fold cross-validation demonstrate that models induced via the presented AI-based pipeline that incorporate machine learning can outperform deep learning and transformer-based models for the prediction of bevacizumab response when the dataset is small.

## 6. Conclusions and Future Work

In this study, we present a novel AI-based computational pipeline to predict the response of ovarian cancer patients to bevacizumab using 250 WSIs obtained from TCIA. First, we resized the original svs images of 54342 × 41048 pixels to average JPEG images of 256 × 256 pixels, and applied HOG and LFDA to construct feature vectors and reduce the dimensionality for a potentially better representation. The new data representation is coupled with labels to train SVR models with several kernels under the regression setting. To assess the generalization capability of the models studied, experimental results utilized five-fold cross-validation. It was demonstrated that our model SVRD+R (our pipeline coupled with the SVR using a radial kernel) outperformed ViT, the best transformer-based model in this study, with a 17% improvement in AUC performance. It also outperformed the best deep learning-based model, MobileNetV2, with a 14.9% improvement in AUC performance. These results demonstrate that our pipeline produces superior results, as well as the feasibility of predicting ovarian cancer drug response using digital pathology images.

In the future, we aim to (1) employ our AI-based computational pipeline to predict drug response pertaining to different cancer types using digital pathology, and (2) develop multimodal deep learning approaches that combine digital pathology images with genomic and other profiling data to improve prediction performance.

## Author Contributions

A.A.: methodology, software, visualization, data curation, and writing— original draft preparation. T.T.: conceptualization, methodology, software, data curation, investigation, supervision, and writing—reviewing and editing. Y.-h.T.: validation and writing—reviewing and editing. All authors have read and agreed to the published version of the manuscript.

## Data Availability Statement

Data are obtained from the cancer imaging archive at https://www.cancerimagingarchive.net/collection/ovarian-bevacizumab-response/ [38,39].

